# Heightened Distraction under Competition in Obsessive-Compulsive Disorder

**DOI:** 10.64898/2026.03.15.711932

**Authors:** Katherine J. McCain, Estelle Ayomen, Arash Mirifar, Heather Simpson Martin, Dominic Demeterfi, Daniel J. McNeil, Gian DePamphilis, Rami Hatem, Robyn Nelson, Grace Melville, Emma Hammes, Alexandra Lee, Ryan McCarty, Mindy Le, Catherine Paciotti, Patricia Coutinho, Carol A. Mathews, Andreas Keil

## Abstract

The present study examine the extent to which attentional resources are allocated toward distracting affective and disorder-relevant pictures under task-relevant competition in participants with obsessive-compulsive disorder (OCD; N = 33) and controls (N = 31). Competition between cues was examined using a foreground task where participants detected coherent motion in a flickering random dot kinematogram (RDK) overlaid on pictures ranging in emotional content (pleasant, neutral, unpleasant, and OCD-evoking pictures). Steady-state visual evoked potentials (ssVEPs) were measured in response to the flickering RDK and served as an index of visuocortical engagement with task-relevant cues. Data were also fitted to the distraction under competition model (DUC), a computational framework of attention selection. Group differences emerged with stronger visuocortical distraction (attenuated task engagement) in the OCD group, driven largely by the unpleasant pictures, followed by the OCD-evoking and pleasant pictures. Furthermore, the DUC model fit well in both groups and supported the results of the univariate analysis, demonstrating the magnitude of the visuocortical distraction observed in response to the unpleasant pictures, and the presence of substantial distraction in response to the OCD-evoking pictures in the OCD group. The present findings provide visuocortical evidence of heightened distraction in response to unpleasant and OCD-evoking pictures under task-relevant competition in OCD.

## Introduction

Exhibiting a lifetime prevalence of approximately 2% (Fawcett et al., 2020), obsessive-compulsive disorder (OCD) is characterized by experiencing obsessive, unwanted thoughts and completing compulsions to alleviate feelings of dread, shame, or discomfort associated with the obsession (Abramowitz et al., 2009; Lack, 2012; Veale & Roberts, 2014). Both obsessions and compulsions can become time-consuming and ritualistic, which can be highly distressing for individuals, increasing the saliency of obsessional contexts and the diversion from goal-driven behavior (Eisen et al., 2006; Moritz, 2008). To this end, early visual perception is often heightened in individuals with OCD and may be indicative of hypervigilance (Chapman et al., 2023; Pallanti et al., 2009; Thomas et al., 2013; Zhang et al., 2017). In addition, evidence of altered visual attentional processing in OCD has emerged across numerous investigations using cognitive and behavioral tasks with neutral, emotional, and disorder-relevant stimuli (for a review, see Ayomen et al., 2025). However, no investigation to date has examined the extent to which task-irrelevant affective and OCD-evoking cues compete with task-relevant cues for limited capacity attentional resources in individuals with OCD. The identification of robust, objective markers of altered visual attention implementation may contribute to our understanding of the neurocognitive basis of OCD and the development of visual domain specific indices of obsessive-compulsive symptoms (OC-symptoms).

Heightened visual perception is often observed in electrophysiological investigations examining event-related visual evoked potential (ERPs) responses in OCD (for a review, see Ayomen et al., 2025). For instance, individuals with OCD showed enhanced P1 (Chapman et al., 2023) and N2 (Lei et al., 2015; Riesel et al., 2017) amplitudes in response to neutral cues in paradigms aimed to capture evidence of altered early visual processing and response inhibition, respectively. In addition, participants with OCD consistently exhibit attentional biases toward aversive or threat-related visual information (for a review, see Cooper & Dunsmoor, 2021), often indicating the presence of altered fear extinction and overgeneralized threat responses (Apergis-Schoute et al., 2017; Cooper & Dunsmoor, 2021; Elsner et al., 2022; Milad et al., 2013). Visuocortical responses to disorder-relevant cues have also been examined in OCD and show enhanced late positive potential (LPP) amplitudes, a measure of sustained attention toward affective cues (Hajcak et al., 2012), in response to disorder-relevant pictures (Paul et al., 2016). However, in the same study, Paul and colleagues (2016) found LPP amplitudes in response to aversive pictures to be larger compared to the amplitude in response to disorder-relevant pictures within OCD participants. Heightened visual processing, particularly in response to aversive or threat-related contexts, has emerged as an index of dysfunction associated with obsessive-compulsive psychopathology. However, it has yet to be examined how this dysfunction manifests under competition.

Affective overlapping interstimulus competition paradigms have been used to examine visuocortical distraction resulting from attentional biases toward affective pictures (Bradley et al., 2012; De Echegaray et al., 2024; Müller et al., 2026). The implementation of this selective attention results in the allocation of attentional resources toward competing sensory inputs, enhancing visual processing of certain inputs while suppressing others (Luck & Gold, 2008). For instance, characteristics of competing stimuli, including their emotional or motivational relevance, can facilitate or bias visual processing, often at the expense of neutral stimuli (Bradley et al., 2012; Desimone & Duncan, 1995; Yiend, 2010). Therefore, the present task was adapted from foundational work by Müller and colleagues (2008) who leveraged interstimulus competition to quantify the attentional trade-off between task-relevant, goal directed cues and emotional task-irrelevant distractor pictures. The authors developed a distraction under competition paradigm which included a foreground change detection task overlaid on naturalistic pictures ranging in emotional valence. The foreground task was flickered at a rate of 8.57 Hz, which allowed for the measurement of steady-state visual evoked potentials (ssVEPs) that served as an index of visuocortical engagement with task-relevant cues. The authors demonstrated that under competition, affective distractor pictures captured attentional resources at the expense of neutral task-relevant cues. Providing a continuous measure of visuocortical engagement, the ssVEP technique is particularly well suited to examine competitive interactions in primary visual cortex in non-clinical and clinical samples (De Echegaray et al., 2024; Deweese et al., 2014; Müller et al., 2008, 2026; Wieser et al., 2011, 2012, 2016).

Other investigations have used similar distraction under competition paradigms and found sustained visuocortical competition in response to affective distractor pictures in clinical samples. For instance, Wieser and colleagues (2012) found evidence of increased visuocortical competition effects in participants with high social anxiety in response to angry faces, compared to pleasant and neutral faces. These competition effects were detected early into the trial (500 ms to 1000 ms) indicating the presence of initial attention capture demonstrated in Müller et al. (2008). Furthermore, following initial attention capture, a sustained period of attention selection toward the task-irrelevant angry faces persisted for another two seconds. Deweese and colleagues (2014) also examined competitive interactions between task cues and task-irrelevant distractor pictures ranging in emotional content and disorder relevance, but in individuals endorsing snake phobia. Participants exhibiting high snake phobia symptoms showed sustained competition in response to the snake pictures, when compared to disorder-irrelevant unpleasant pictures, extending several seconds after picture onset. Critically, in both cases, this pattern of sustained competition was specific to the presence of disorder-relevant cues with a negative emotional valence.

The present study leveraged ssVEPs to quantitatively and continuously measure attentional allocation between task-relevant cues and task-irrelevant distractors in individuals with OCD and controls. A taxing coherent motion detection task (random dot kinematogram; RDK) was overlaid on naturalistic distractor pictures ranging in emotional content, including disorder-relevant (OCD-evoking), pleasant, unpleasant, and neutral pictures. Resembling the OCD-evoking picture themes identified by Paul and colleagues (2016), the present study included OCD-evoking pictures covering several OCD symptom categories. Disorder-relevant images were chosen to primarily elicit OCD symptoms in participants who may have endorsed obsessions or compulsions related to aggressive, contamination, cleaning/washing, checking, symmetry, or counting. Given that obsessions and compulsions, across individuals, are more idiosyncratic than fears in specific phobia and social anxiety disorder (Broekhuizen et al., 2023; Brooks et al., 2018; Moritz et al., 2009), evoking attention biases in participants with OCD warrants the inclusion of multiple symptom-provoking dimensions. However, some pictures may be neutral to many observers, because they include content that, for example, is related to symmetry, checking, or counting, not known to elicit aversive responses.

Due to the salient nature of affective naturalistic scenes (Wöstmann et al., 2022), we expected decreased visuocortical engagement or attenuation of the task-evoked ssVEP envelope amplitudes following affective distractor picture onset, relative to neutral picture onset, regardless of group. Furthermore, we expected to observe group differences in distraction under competition and hypothesized that these differences would be driven by specific picture conditions. For instance, the pleasant, unpleasant, and OCD-evoking picture conditions were expected to prompt greater visuocortical distraction under competition in the OCD group compared to the control group. We also anticipated poorer task performance on trials where unpleasant, pleasant, and disorder-relevant distractors were presented for the OCD group compared to the control group. To further elucidate the attentional trade-off between task-relevant cues and task-irrelevant affective and disorder-relevant pictures, the present study also quantified visuocortical distraction by fitting the data to the distraction under competition model (DUC; De Echegaray et al., 2024; Müller et al., 2026), a framework for selective attention. The DUC model characterizes competitive interactions between task-evoked and distractor-evoked responses in visual cortex (De Echegaray et al., 2024; Müller et al., 2026), producing meaningful parameter estimates of visuocortical distraction. We expected the DUC model to fit the empirical data, corroborate an attentional shift from task cues to affective and disorder-relevant distractors, and indicate increased visuocortical distraction in the OCD group compared to the control group.

## Methods

### Participants

Thirty-three participants with OCD (27.42 ± 9.35 years) and 31 control participants (38.13 ± 18.97) were included in the present study and recruited from psychiatry and psychology clinics at the University of Florida and through flyers placed in the local community. Supplemental Table 1 shows the distribution of demographic data across groups, including gender, race, ethnicity, and age. Sample size determinations were based on *a priori* statistical power (1-β) using G*Power software (Faul et al., 2007) and included the following specifications: Effect size f = 0.20; α = 0.05; (1-β) = 0.9; correlation among repeated measurements = 0.5; nonsphericity correction e = 1; test family = F test, and statistical test = ANOVA repeated measures, within-between interaction. The power analysis showed that at a minimum, 56 participants were required to detect low to medium effects, so the present study is well-powered. All activities related to this research project were completed as a part of a large-scale research initiative investigating obsessive-compulsive anxiety spectrum disorders and approved by the University of Florida International Review Board in accordance with the Declaration of Helsinki.

### Clinical assessments

Participants were assessed using a battery of self-report questionnaires and underwent a semi-structured clinical interview conducted by a clinician trained in research procedures. OCD was assessed using the Yale-Brown Obsessive Compulsive Scale (Y-BOCS; Goodman et al., 1989) the Obsessive-Compulsive Inventory (OCI-R; Foa et al., 2002), and the Mini Neuropsychiatric Interview (MINI; Sheehan et al., 1998). The Structured Interview for Hoarding Disorder (SIHD; Nordsletten et al., 2013) and Saving Inventory-revised (SI-R; Frost et al., 2004) was used to assess for hoarding disorder. The SNAP-IV (Bussing et al., 2008) was used to assess for Attention Deficit and Hyperactivity Disorder (ADHD). Medical records, when available, were obtained to provide additional clinical information. A best estimate process was used to assign DSM-5 diagnoses. Best estimators were blinded to the case/control status of the participant and reviewed all available clinical data (i.e., self-report questionnaires, clinical interviews, and medical records; Nutley et al., 2020) for a given participant. Assignments of “definite,” “probable,” “not present,” or “not determined” (e.g, not enough data were present to make a determination) were made for each DSM-5-TR diagnosis. In the event of probable diagnoses, two best estimates were independently completed, and a subsequent consensus diagnosis was reached. In the event that a consensus could not be reached between two best estimators, a third best estimator was assigned.

### Inclusion and exclusion criteria

Participants were included in the study if they were 18 years of age or older. OCD participants were required to meet criteria for a lifetime history of OCD as defined by the DSM-5. Control participants were included if they did not meet criteria for OCD. Those with psychosis, known intellectual disability, head trauma with loss of consciousness, or medical or neurological conditions known or suspected to affect cognitive function were also excluded. A lifetime or current history of other psychiatric diagnoses were not exclusionary in either group. Participants with fewer than 50% of trials remaining in any condition after EEG artifact rejection, were excluded from the analysis. Finally, participants exhibiting less than 50% task accuracy across all picture conditions were excluded from the analysis.

### Stimuli and procedure

Visual stimuli were created in Psychtoolbox for Matlab (version 3.0.18) and presented on a Samsung LS23A950 Monitor with a 120 Hz refresh rate and resolution of 1920 x 1080. The distraction under competition paradigm included a foreground coherent motion detection task overlaid on naturalistic distractor pictures ranging in emotional content and disorder relevance. The motion stimulus included 150 bright yellow dots (0.29° of visual angle; see Figure 1) constrained by a central circular region of the screen (9.20° of visual angle) and maintained constant motion while flickering at a frequency of 8.57 Hz. Pleasant, neutral, and unpleasant distractor pictures were chosen from the International Affective Picture Set (IAPS; Lang et al. 1997), and OCD-evoking pictures were carefully curated from a database of openly licensed pictures. OCD-evoking pictures were chosen to primarily elicit OCD symptoms in participants who endorsed obsessions or compulsions related to aggressive, contamination, cleaning/washing, checking, symmetry, or counting (Goodman et al., 1989).

**Figure 1.**
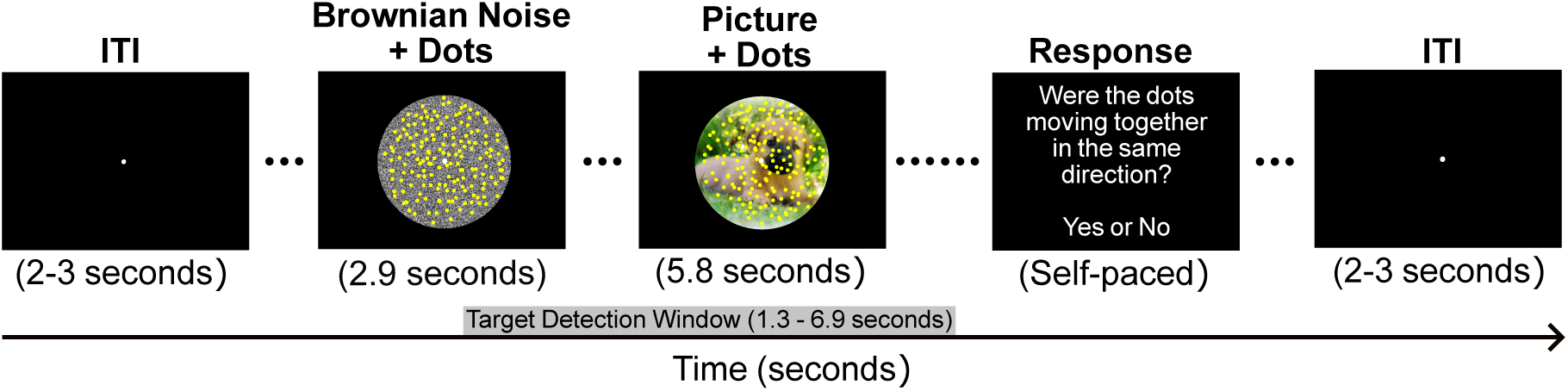
Distraction under competition task. Trials began with a period of Brownian noise (2.90 s). The distraction under competition period (5.80 s) was marked by the onset of the distractor pictures subtending the same annulus defined in the Brownian noise period combined with the continuation of the motion stimulus overlayed. On target trials half of the dots moved together in coherent motion in one direction (45°, 135°, 225°, or 315°) once per trial (target detection window). On non-target trials, the dots never moved in the same direction. Following each trial, participants were asked to indicate whether they detected coherent motion using the mouse to select ‘yes’ or ‘no’ during the self-paced response period.

Trials began with a period of Brownian noise (2.90 s) where the motion stimulus was superimposed over a central circular background with the RBG value of each pixel ranging randomly between 0-255, which created the perception of a gray-scale background (9.20° of visual angle; see Figure 1). The Brownian noise period functioned to attenuate transient visuocortical responses to luminance changes at the onset of the distractor pictures behind the motion stimulus (Deweese et al., 2014). The distraction under competition period (5.80 s) was marked by the onset of the distractor pictures subtending the same annulus defined in the Brownian noise period combined with the continuation of the motion stimulus overlaid (see Figure 1). Throughout the trial, the motion stimulus was presented for a total of 8.70 seconds, and participants were asked to maintain fixation by directing their gaze to a white fixation dot in the center of the screen (0.34° of visual angle; see Figure 1). On target trials, half of the dots could move together in coherent motion in one direction (45°, 135°, 225°, or 315°) once per trial, and this coherent motion could only occur between 1.3 and 6.9 seconds (coherent motion detection time window) into the trial. On non-target trials, the dots never moved in the same direction, maintaining constant random motion throughout. The task consisted of 30 trials in each of the four picture conditions. Target and non-target trials were evenly distributed, so all picture conditions included 15 target and 15 non-target trials that were then randomized within the conditions. Following each trial, participants were asked to indicate whether they detected coherent motion using the mouse to select ‘yes’ or ‘no’ during the self-paced response period. After the self-paced response period, the inter-trial interval ranged from 2-3 seconds. The task began with a brief practice, and upon completion of the entire experimental task, participants provided subjective valence and arousal ratings for each picture (see Figure 2) using the 9-point Self-Assessment Manikin scale (SAM; Bradley & Lang, 1994). For valence, a rating of 1 indicated negative valence (unpleasant) and a rating of 9 indicated positive valence (pleasant). For arousal, a rating of 1 indicated low arousal (evoking a feeling of calm) and a rating of 9 indicated high arousal (evoking excitement).

**Figure 2.**
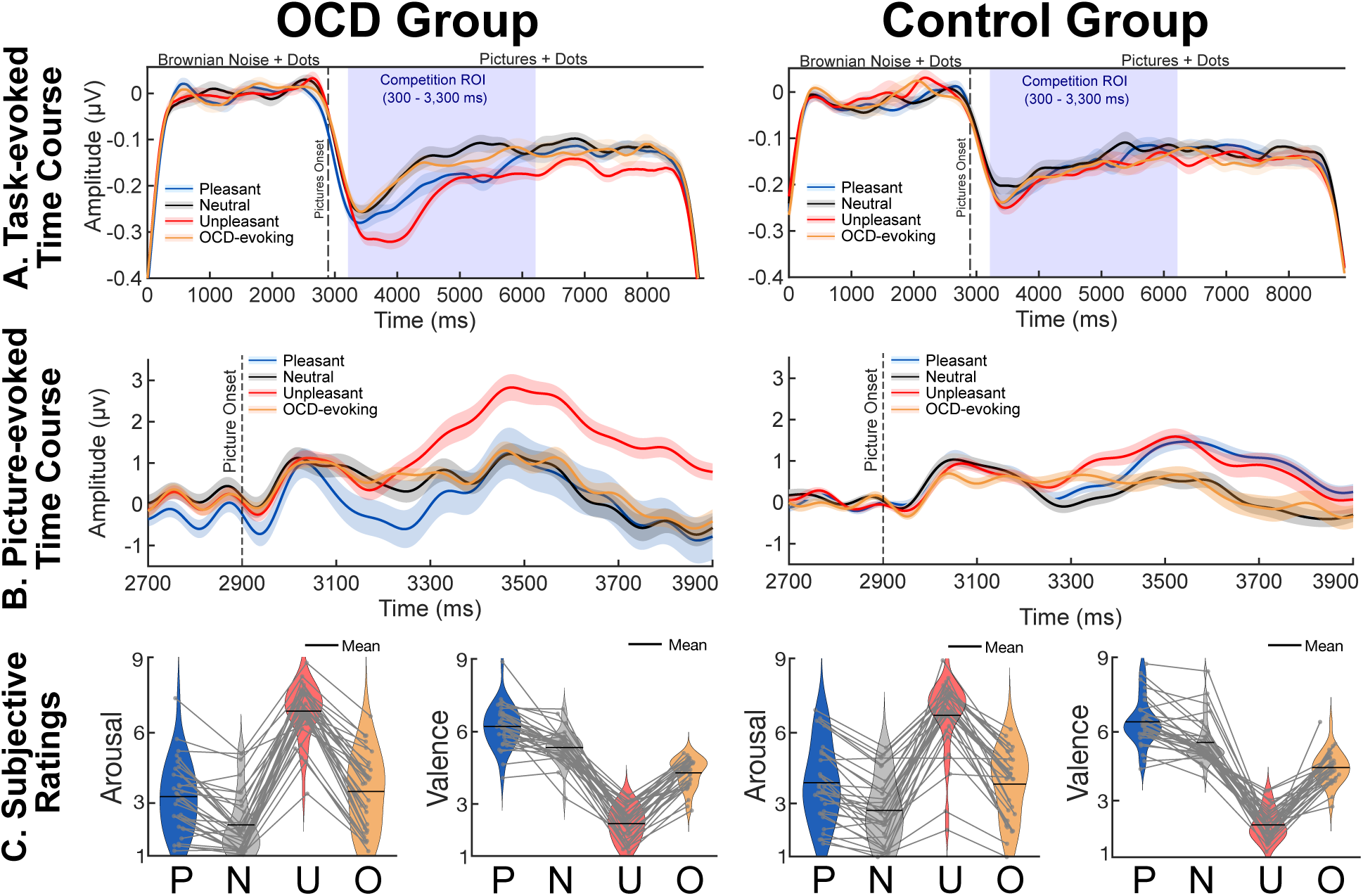
(A) Grand average task-evoked ssVEP envelopes were computed across the sensors contributing to the interaction effect identified in the cluster-based permutation analysis. The shaded region around the solid lines indicates within-subject error. (B) Picture-evoked time course included in the distraction under competition (DUC) model. The shaded region around the solid lines indicates within-subject error. (C) Subjective valence and arousal ratings across conditions (pleasant, P; neutral, N; unpleasant, U; OCD-evoking, O) using the self-assessment manikin (SAM). For valence, a rating of 1 indicated negative valence (unpleasant) and a rating of 9 indicated positive valence (pleasant). For arousal, a rating of 1 indicated low arousal (evoking a feeling of calm) and a rating of 9 indicated high arousal (evoking excitement). Light gray dots and the lines connecting them indicated individual subject average arousal and valence ratings for each condition.

**Figure 3.**
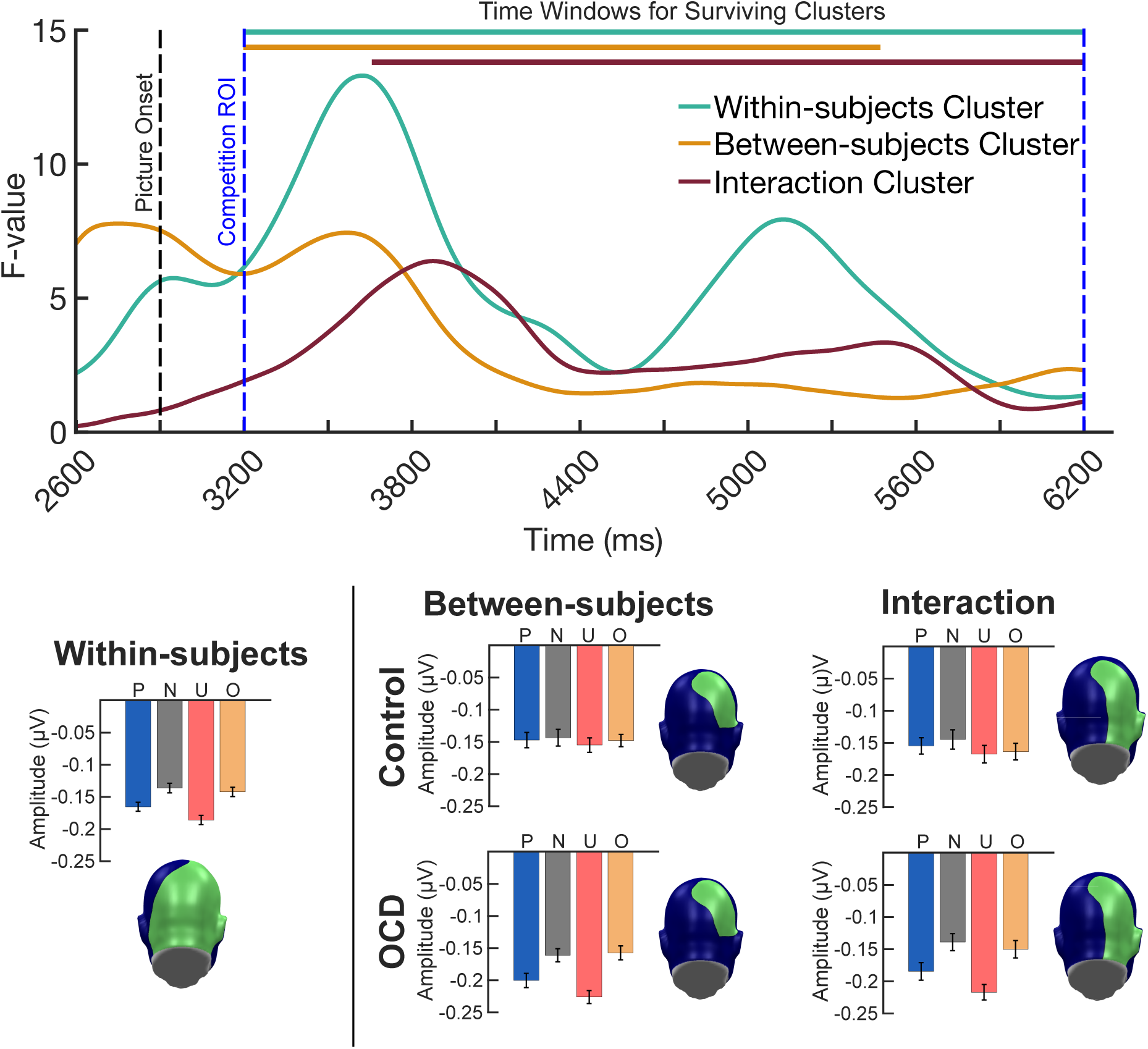
Evidence of group differences exhibiting stronger competition effects for the OCD group, driven largely by the unpleasant pictures, and less so by the OCD-evoking pictures. Time courses were obtained by averaging the resulting F-values from the mass univariate mixed-factor planned contrast, across the sensors identified in each of the clusters that survived cluster-based permutation analysis. Colored bars at the top of the time course indicated the length of the time window identified for each cluster that survived cluster-based permutation analysis. Topographies indicate the sensors identified in the within-subjects, between-subjects, and interaction clusters that survived cluster-based permutation analysis. Average task-evoked envelope amplitudes for each condition conditions (pleasant, P; neutral, N; unpleasant, U; OCD-evoking, O) were computed across the within-subjects, between-subjects, and interaction cluster time window and sensors are shown as bar plots.

### EEG data acquisition

Participants were seated in an electrically shielded recording chamber 100 cm from the stimulus presentation monitor. Continuous EEG data was recorded using an electrical geodesics (EGI) high-density, high impedance 257 channel system. Online recording parameters included sensor Cz as the online reference and a sampling rate of 1000 Hz. In addition, impedance levels were kept below 60 kΩ throughout the recording.

### EEG data processing

Offline, the continuous EEG data were processed using custom automated preprocessing and postprocessing pipelines created using the EEGLAB MATLAB toolbox (Delorme & Makeig, 2004) and are openly available in the project repository (https://osf.io/s73d5/overview).

#### Task-evoked responses

##### Preprocessing

The preprocessing pipeline began by down sampling the continuous EEG data from 1000 Hz to 500 Hz. Next, a 3 Hz high-pass Butterworth filter (4^th^ order) and a 30 Hz low-pass Butterworth filter (9^th^ order) were applied to the continuous data. Ocular artifact correction was performed using the open-source BioSig package (BioSig, 2005) and an automated correction procedure (Schlögl et al., 2007). The continuous EEG data were then segmented 600 ms prior to motion stimulus onset and 9000 ms post-stimulus onset to capture both the Brownian noise and distraction under competition periods. Next, artifact correction and rejection procedures were applied to the segmented data using a robust procedure developed for data acquired through high density EEG systems. The Statistical Control of Artifacts in Dense Sensors Arrays (SCADS; Junghöfer et al., 2000) computes a compound data quality index for each participant at the individual channel and trial level. The compound data quality index is comprised of three statistical parameters with set thresholds including absolute mean amplitude, standard deviation, and maximum transient voltage change. Channel activity throughout the entire recording (global level) and at the single trial level was flagged for data quality when the quality index exceeded 2.5 standard deviations above the median index for that channel or trial, respectively. Flagged channels at the global and single trial level were interpolated by means of a 2D spline interpolation in which estimations of the values for the flagged channels were based on all channels. After interpolation, individual trials were flagged and rejected when the data quality index of that trial exceeded 1.25 times the median data quality index. After artifact rejection and correction, individual subject grand means were obtained by averaging surviving artifact free trials across the distractor picture conditions.

##### Postprocessing

The postprocessing pipeline functioned to transform the individual subject picture condition averages into time-varying estimates of the phase and amplitude of the steady-state evoked potential response to the flickering motion stimulus. To prepare for the transform, individual subject picture condition averages were referenced to the average reference and narrowly filtered using a band-pass Butterworth filter (9th order) with a lower frequency cut-off of 8.07 Hz and upper frequency cut-off of 9.07 Hz. The purpose of the narrow band-pass filter was to isolate the frequency band of interest, extracting the ssVEP response to the motion stimulus at 8.57 Hz. Next, the narrowly band-passed picture condition averages were Hilbert transformed using the FreqTag MATLAB toolbox (Figueira et al., 2022). Time-varying estimates of the amplitude and phase of the ssVEP response to the motion stimulus were extracted using the Hilbert transform to create an envelope of the signal across time and averaged across picture conditions. The resulting individual subject grand mean Hilbert envelopes provided continuous measures of visuocortical engagement with the task-relevant cues in visual cortex (Wieser et al., 2016). Individual subject grand mean responses were baseline adjusted to the average voltage of -2500 to -400 ms prior to picture onset and included in both the mass univariate analysis approach and DUC model. The baseline adjustment time window extended throughout much of the Brownian noise period (excluding the first and last 400 ms to attenuate edge artifacts) to subtract the baseline ssVEP response to the motion stimulus in the absence of the competitive interaction (see Figure 2).

#### DUC model

##### Preprocessing

The preprocessing pipeline for the picture-evoked responses used for DUC model fitting was virtually identical to the ssVEP preprocessing pipeline with the exception of the high- and low-pass filter parameters. The continuous EEG data were filtered using a 0.1 Hz high-pass Butterworth filter (4th order) and an 8 Hz low-pass Butterworth filter (10th order). The purpose of the strict low-pass filter was to extract the majority of the transient event-related potential response from the oscillatory steady-state evoked potential response to obtain competing visual responses to the affective distractor pictures and task-relevant motion stimulus, respectively across the same time course. This method has been applied in previous studies conducted by Müller & Hillyard (2000) and Schönwald & Müller (2014), demonstrating the feasibility of isolating transient stimulus-locked responses from steady-state signals using a low-pass filter with a cut-off frequency lower than the stimulation rate. Furthermore, the stimulation rate of 8.57 Hz guards against any potential overlap in the picture-evoked response and the task-evoked response (Bekhtereva et al., 2018). Individual subject grand means were baseline correct to the average voltage 200 ms prior to distractor picture onset (see Figure 2).

##### Postprocessing

Additional data preparation steps were taken to further isolate the distractor-evoked response from the task-evoked oscillatory signal and to standardize the two visual evoked response streams for optimal parametrization. The distractor-evoked signal was averaged across an *a priori* parietal-occipital cluster of 11 sensors surrounding electrode Pz (80, 89, 90, 100, 101 (Pz), 110, 118, 127, 128, 129, 130, 131), often included in analyses of sustained picture-evoked responses toward affective visual stimuli (Cuthbert et al., 2000; Hajcak et al., 2012; Keil et al., 2002; Schönwald & Müller, 2014; Weinberg & Hajcak, 2010). The task-evoked response was averaged across sensor Oz. To account for the stimulus-locked nature of the picture-evoked signal, a duration of 1000 ms after distractor picture onset was included in the DUC model analysis for both the picture- and task-evoked responses. Both a 100-point centered moving average and 50 ms cosine window were applied to the individual subject picture-evoked time course across groups. Additional baseline correction was applied to the picture-evoked time course. The resulting task and picture-evoked individual subject grand mean time courses were Z-transformed across groups and included in the DUC model.

### Statistical analyses

#### Mass univariate approach

##### Task-evoked responses

A mass univariate approach allowed for the examination of the ssVEP responses without the need for rigid *a priori* sensor and time window determinations (Groppe et al., 2011). This approach is particularly well-suited for the statistical analysis of ssVEPs recorded using a high-density EEG system because of the increased spatial and temporal capabilities afforded with this technique (Groppe et al., 2011). The present analysis examined the emergence of visuocortical distraction across groups and picture conditions at each sensor and time point in the task-evoked response (Hilbert envelopes) to the motion stimulus after the distractor picture onset. We hypothesized that distraction would differ across groups, and that these differences would be driven by the affective and OCD-evoking picture conditions. To this end, we directly tested this hypothesis using mixed-factor planned contrasts with group-specific within-subject weights, placing greater weight on the affective and OCD-evoking picture conditions for the OCD group compared to the control group. To ensure robustness of the findings, the mixed model analyses were also repeated with standard weights.

Reflective of the hypotheses as described in the introduction, the planned contrast weights for the control group were -1 (Pleasant), 1 (Neutral), -1 (Unpleasant), 1 (OCD-evoking) and the OCD group weights were -0.5 (Pleasant), 2 (Neutral), -1 (Unpleasant), -0.5 (OCD-evoking). The control group weights (standard arousal weights) were modeled after the characteristic, reliable pattern of electrocortical activation, often observed in nonclinical samples, where affective IAPS pictures captured greater attentional resources compared to neutral pictures (Bradley et al., 2012; Hindi Attar et al., 2010; Lang & Bradley, 2010; Müller et al., 2008; Schönwald & Müller, 2014). Moreover, we expected the responses to the OCD-evoking pictures to mirror the responses to neutral stimuli in the control group, so the contrast weights were equivalent. Conversely, contrast weights for the OCD group were distributed so there was greater contrast magnitude, compared to the standard arousal weights, between the affective and OCD-evoking picture conditions and neutral picture condition (symptom-specific weights).

To control for multiple comparisons, a nonparametric cluster-based permutation analysis was used to identify significant effects of distraction, across time and space (Maris & Oostenveld, 2007). The sensitivity of the cluster-based permutation analysis was optimized according to Maris and Oostenveld (2007): Biophysically motivated constraints were placed on the sensors and time points included in the cluster-based permutation analysis to further robustness and emphasize physiologically plausible results. For instance, we expected to detect task-evoked ssVEP responses across parietal and occipital sensors, given that visual cortex primarily contributes to the emergence of ssVEP responses (Di Russo et al., 2006). Furthermore, studies employing variations of distraction under competition paradigms in anxious samples have shown reliable individual differences in competition emerging several hundred milliseconds after affective picture onset and persisting for several seconds (Deweese et al., 2014; Wieser et al., 2012). Therefore, occipital-parietal sensors and time-points spanning a 3 second *a priori* competition window of interest beginning 300 ms after picture onset were included in the permutation analysis. Statistically significant clusters were identified via a conservative F_max_ procedure, increasing our ability to reliably detect large effects (Tebbe et al., 2026). Permuted within-subjects, between-subjects, and interaction F-scores were calculated separately across 1,000 permutations at each sensor and time point included in the analysis. Subjects included in each of the permutations were randomly sampled with replacement and their corresponding picture condition labels were shuffled. Mass univariate and cluster-based permutation analyses were carried out using custom Matlab scripts available in the project repository (https://osf.io/s73d5/overview).

#### Control analysis

A control analysis was carried out to evaluate visuocortical engagement with task-cues in response to the affective and neutral picture conditions, excluding the OCD-evoking picture condition from the planned contrast (three weight contrast: -1, Pleasant; 2, Neutral; -1, Unpleasant). The control analysis was constrained by the same cluster-based permutation-controlled analysis detailed above.

### Modeling visuocortical trade-off driven by attention biases

The DUC model functions to characterize competitive interactions between task-evoked and distractor-evoked responses in visual cortex (De Echegaray et al., 2024; Müller et al., 2026). This is accomplished by modeling feedforward processes, the initial stimulus gain of the distractor, and the subsequent competition for visuocortical representation between task-evoked and distractor-evoked cues (De Echegaray et al., 2024; Müller et al., 2026). Integrating the biased competition (Desimone & Duncan, 1995) and sensory gain amplification (Hillyard et al., 1998) conceptual models of selective attention, the DUC model includes three parameters of interest to model the attentional “push and pull” between competing visual stimuli. The β_D_ parameter models the initial content-selective response to the distractor stimulus, the ϕ parameter models the early inhibitory influence of the distractor cues on the task-relevant cues, and the λ parameter models sustained competition between the distractor and task-evoked response (De Echegaray et al., 2024; Müller et al., 2026).

In an identical procedure to De Echegaray et al. (2024) and Müller et al. (2026), the DUC model was fit to the empirical data using non-linear regression and the Levenberg-Marquart algorithm (nlinfit in MATLAB). 5,000 bootstrapped grand mean task-evoked envelopes and picture-evoked waveforms were estimated to quantify the distribution of the parameters in the sample. Grand average envelope and waveform magnitude was expressed as percent change from baseline, and all model parameters varied freely. Finally, a non-parametric Bayesian bootstrap approach (Ahumada et al., 2025; Efron, 2011) was used to compare the distributions of the parameter estimates within picture conditions and between groups. Mean square error (MSE) was calculated across each distribution and the same Bayesian bootstrap approach was used to compare model fit indices within and across groups. The present application of the DUC model was performed using custom Matlab scripts available in the project repository (https://osf.io/s73d5/overview).

### Behavioral data

Task performance was also analyzed using the same mixed-factor planned contrasts with the group-specific within-subject weights detailed above to examine the effects of picture content (pleasant, neutral, unpleasant, and OCD-evoking) and group (OCD and control) on task accuracy. Task accuracy was quantified as the proportion of correct responses (indicated ‘yes’ when there was a coherent motion event in target trials and ‘no’ when there was no coherent motion on non-target trials) to the total number of responses.

## Results

### Demographics

Demographic data were examined across groups (see Supplemental Table 1) and indicated that the control and OCD groups did not differ across gender (*X*^2^(2*, N =* 64) = 2.59, p = 0.27, Cramer’s V = 0.20), race (*X*^2^(4, *N =* 64) = 7.41, p = 0.12, Cramer’s V = 0.34), or ethnicity *(X*^2^(1, *N =* 64) = 0.27, p = 0.60, Cramer’s V = 0.07). However, participants in the control group (M = 38.13, SD = 18.97) were significantly older (t(43.16) = 2.84, p = .007, *d* = 0.72) compared to the participants in the OCD group (M = 27.42, SD = 9.35 years).

### Trial counts

Repeated measures ANOVAs were carried out to examine the number of trials included in the analyses. There were no significant differences in the number of trials rejected across conditions after application of the ssVEP preprocessing pipeline (F(3, 186) = 0.43, p = 0.73, partial η^2^ = 0.01). However, there was a significant difference in the number of trials rejected due to poor data quality between groups (F(1, 62) = 4.42, p = 0.04, partial η^2^ = 0.07). Descriptively, the average number of ssVEP trials rejected in the control group was about one trial more than the average number of trials rejected in the OCD group, regardless of condition. For the picture-evoked preprocessing pipeline, there were no significant differences in the number of trials rejected between groups (F(1, 62) = 1.29, p = 0.26, partial η^2^ = 0.02) or across conditions (F(3, 186) = 0.70, p = 0.55, partial η^2^ = 0.01). Similarly, the average number of picture-evoked trials rejected in the control group was just under one trial more than the average number of trials rejected in the OCD group, regardless of condition.

### Behavioral data

The results of the mixed-factor planned contrast revealed a significant effect of condition (F(1, 192) = 15.16, p < .001, partial η^2^ = 0.02) on task accuracy. However, the effect of group was not significant (F(1, 192) = 0.02, p = 0.88, partial η^2^ = 0.0001). Regardless of group, these results indicated poorer task performance on trials where affective and disorder-relevant pictures were presented compared to neutral pictures.

### Mass univariate analysis across time and sensors

Cluster-based permutation analysis revealed statistically significant clusters of sensors exhibiting cluster sums (F-value sum for each cluster) greater than the permuted F-maximum (maximum cluster sum of permuted F-values) across time. Within subjects, a large cluster containing 101 occipital-parietal sensors over medial posterior regions exceeded (∑F-value_observed_ = 710,269) the permuted within-subjects F-maximum (∑F-value_perm_ = 26,950) and spanned the entire *a priori* competition time window of interest (3,200 ms to 6,200 ms or 300 ms to 3,300 ms post picture onset). Between subjects, a moderately sized cluster emerged across 27 right lateralized occipital-parietal sensors exhibited a cluster sum (∑F-value_observed_ = 97,221) exceeding the permuted between-subjects F-maximum (∑F-value_perm_ = 14,732) and spanned a considerable amount of the competition time window from 3,200 ms to 5,574 ms. The interaction cluster included 45 right lateralized occipital-parietal sensors and displayed a cluster sum (∑F-value_observed_ = 119,798) surpassing the permuted interaction F-maximum (∑F-value_perm_ = 93,685). Significant individual differences in competition were detected around 760 ms after picture onset and persisted across the remainder of the *a priori* competition time window (3,660 ms - 6,200)^1^.

### Control analysis

In addition, the control analysis revealed a significant main effect of condition, but not of group when the OCD-evoking picture condition was left out of the analysis. The resulting cluster included 97 occipital-parietal sensors, which were subsumed in the within-subjects cluster previously identified in the 8-weight cluster-based permutation analysis. The cluster sum (∑F-value_observed_ = 533,011) exceeded the permuted F-maximum (∑F-value_perm_ = 7,781) and again spanned the entire *a priori* competition time window interest (3,300 ms to 6,300 ms or 300 ms to 3,300 ms post-picture onset)^2^.

### DUC Model Fitting

Table 1 shows the best-fitting parameter estimates by condition for each of the three parameters of interest. Also included in Table 1 are the resulting log base 10 Bayes factors (Log10BF) computed using a Bayesian bootstrap approach (Ahumada et al., 2025; Efron, 2011) to compare the bootstrapped distributions of the parameter estimates across conditions. Bayes factors were interpreted according to Jeffreys (1961), where log10BF values greater than |1| were considered strong evidence, values greater than |1.5| were considered very strong evidence, and values greater than |2| were considered decisive evidence of an effect. Bayesian comparisons across picture conditions resulted in an overall trend of increased affective picture interference (enhanced neural responses toward emotional stimuli compared to neutral stimuli; Bradley et al., 2012) in the pleasant and unpleasant picture conditions for the control group, and the unpleasant picture condition for the OCD group. For the OCD group, the content-selective parameter comparison across the unpleasant and neutral picture conditions revealed decisive evidence of an increased initial content-selective response toward the unpleasant pictures. For the control group, the content-selective parameter comparison indicated strong to very strong evidence of an increased initial content-selective response for the unpleasant and pleasant pictures, respectively. Table 2 shows the Bayes factors (Log10BF) obtained for each comparison of the parameter estimates across groups and picture conditions. Overall, comparisons across groups confirmed the strength of the attentional impact of the unpleasant pictures in the OCD group, supporting the results of the mass univariate analysis. Comparisons also indicated that the OCD-evoking pictures substantially impacted attentional engagement with the task-relevant cues in the OCD group compared to the control group, not readily observed in the univariate analysis.

**Table 1.** Best fitting DUC model parameter estimates of the three parameters of interest can be seen in the first four columns (negative parameter estimates indicate greater distraction). In addition, comparisons of the parameter estimate distributions by means of a Bayesian bootstrap procedure can be seen in the last four columns. Bootstrap distributions were compared within groups: neutral (*θ_1_*, N) compared to pleasant (*θ_2_*, P), neutral (*θ_1_*, N) compared to unpleasant (*θ_2_*, U), pleasant (*θ_1_*, P) compared to unpleasant (*θ_2_*, U), and neutral (*θ_1_*, N) compared to OCD-evoking (*θ_2_*, O) picture conditions. For the comparisons between affective and neutral picture conditions, greater content-selective initial response (β_D_), early distractor inhibitory influence (ϕ), or late competition (λ) in response to affective and disorder-relevant pictures (increased distraction) was indicated by positive log base 10 Bayes factors (log10BF) and negative log10BF values indicated increased β_D_, ϕ, or λ in response to the neutral pictures (decreased distraction). For the pleasant (θ*_1_*, P) and unpleasant (θ*_2_*, U) picture condition comparison, positive log10BF values indicated increased β_D_, ϕ, or λ in response to the unpleasant pictures, and negative log10BF values indicated increased β_D_, ϕ, or λ in response to the pleasant pictures. Log10BF values were interpreted according to Jeffreys (1961) scale where values greater than 1 indicated strong evidence of distraction under competition.

| Group | Parameter | Pleasant | Neutral | Unpleasant | OCD | Log10BF<sub>NP</sub> | Log10BF<sub>NU</sub> | Log10BF<sub>PU</sub> | Log10BF<sub>NO</sub> |
| --- | --- | --- | --- | --- | --- | --- | --- | --- | --- |
| Control | Content-selective picture response ($\beta_D$) | -14.55 | 1.83 | -3.41 | 0.73 | 1.84 | 1.36 | -0.60 | 0.31 |
| | Early inhibitory influence ($\phi$) | -0.27 | 3.68 | -0.94 | 3.17 | 0.75 | 0.71 | 0.36 | 0.28 |
| | Late competition ($\lambda$) | 0.35 | -0.48 | -1.42 | -1.41 | -0.09 | 0.26 | 0.41 | 0.14 |
| OCD | Content-selective picture response ($\beta_D$) | -2.16 | -0.52 | -9.43 | -0.75 | 0.30 | 2.99 | 1.26 | 0.07 |
| | Early inhibitory influence ($\phi$) | -0.94 | -4.50 | -0.81 | -4.75 | -0.54 | -0.57 | 0.20 | 0.07 |
| | Late competition ($\lambda$) | -1.40 | -1.85 | -1.92 | -3.69 | -0.11 | 0.02 | 0.14 | 0.41 |

**Table 2.**
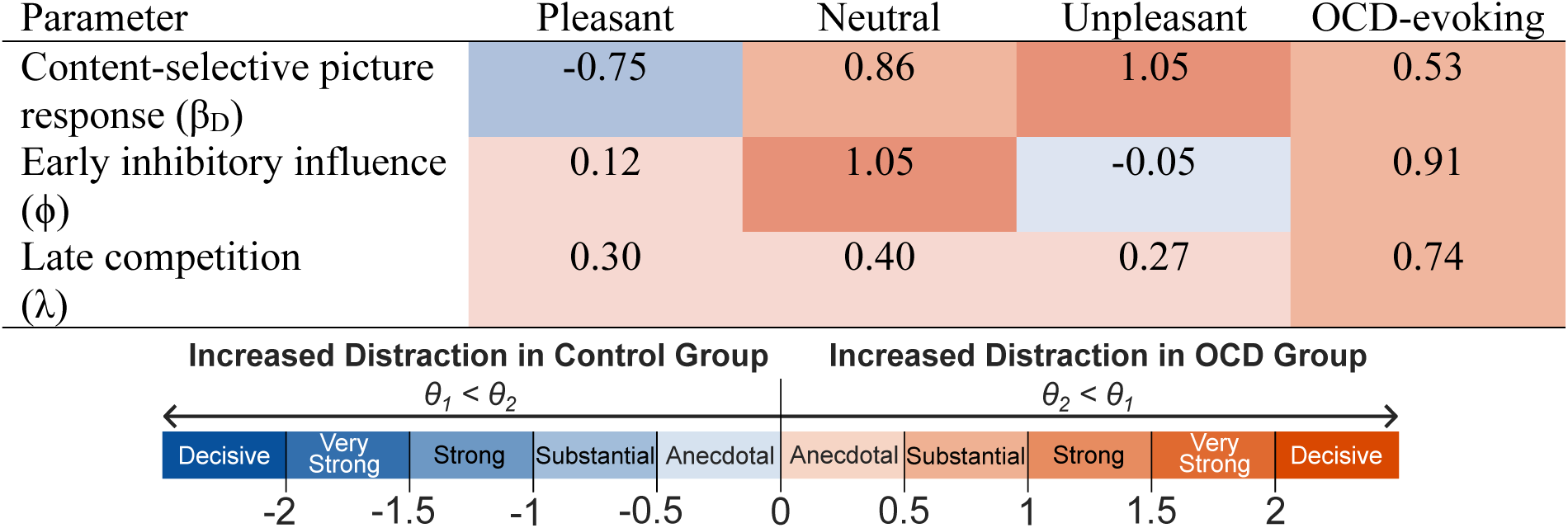
Group comparison of parameter estimates bootstrap distributions using the Bayesian bootstrap procedure (Log10BF) and interpretation reported in Table 1. Bootstrap distributions of parameter estimates obtained for the control group (*θ_1_*) were compared to the OCD group (*θ_2_*). Positive log base 10 Bayes factors (log10BF) indicated increased content-selective initial response (β_D_), early distractor inhibitory influence (ϕ), or late competition (λ) in the OCD group. Negative log base 10 Bayes factors (log10BF) indicated increased content-selective initial response (β_D_), early distractor inhibitory influence (ϕ), or late competition (λ) in the control group.

The DUC model exhibited good fit with the empirical data (see Figure 4). Model fit within groups was assessed using average mean square error (mMSE) computed across the distributions of the MSE values obtained for each resample. Model fit across groups was examined using the same Bayesian bootstrap approach where distributions of MSE values for each condition were compared across groups. Positive log10BF values indicated greater model fit for the OCD group and negative log10BF values indicated greater model fit for the control group. Bayes comparisons showed strong to very strong evidence of greater model fit in the OCD group for the neutral (log10BF = 1.41) and pleasant (log10BF = 1.95) picture conditions, respectively. Comparisons showed similarly strong evidence of greater model fit in the control group for the unpleasant (log10BF = -1.32) picture condition. For the OCD-evoking condition, Bayes comparisons showed anecdotal evidence of greater model fit in the OCD group for the OCD-evoking (log10BF = .44) picture condition.

**Figure 4.**
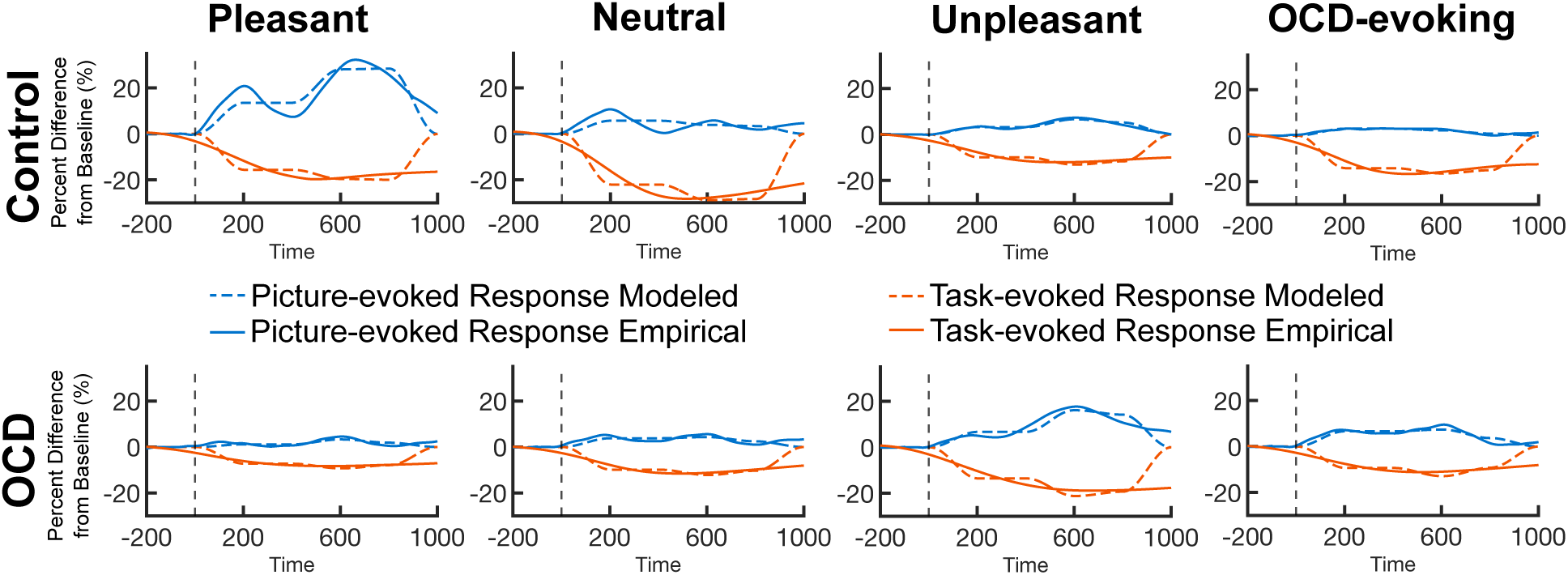
Best-fitting picture-evoked and task-evoked modeled responses fit to empirical picture-evoked and task-evoked responses.

## Discussion

The present study examined the extent to which task-relevant and task-irrelevant affective and OCD-evoking pictures compete for limited capacity attentional resources in individuals with OCD and controls. In line with our first hypothesis, task-evoked ssVEP envelope amplitudes were attenuated upon presentation of the distractor pictures, more so for affective compared to neutral pictures, regardless of group. The differential impact of affective picture content on task-evoked ssVEP envelope amplitudes aligns with accumulating evidence of enhanced visual processing of emotional pictures (pleasant and unpleasant IAPS pictures), often at the cost of concurrently presented neutral task-relevant cues (Bradley et al., 2012; Deweese et al., 2016; Hindi Attar et al., 2010; Keil et al., 2005). Group differences in distraction under competition were, driven primarily by the unpleasant distractor pictures, followed by pleasant and OCD-evoking pictures. The OCD group exhibited increased distraction in the affective and disorder-relevant conditions compared to the control group. This interaction began around 760 ms after affective picture onset, in a right lateralized occipital-parietal cluster of sensors and persisted throughout the *a priori* competition window. The lateralization of the interaction effect is not surprising in the light of past research: Functional magnetic resonance imaging (fMRI) has shown increased bilateral BOLD activation in the occipital gyrus, and right lateralized BOLD activation in the inferior and superior parietal lobes in response to emotional, compared to neutral pictures (Lang et al., 1998). In addition, the time course of the interaction effect was consistent with both the onset and duration of the group differences in competition identified by Wieser and colleagues (2012) and Deweese and colleagues (2014) in anxious samples.

The interaction effect described above provides evidence of altered attentional implementation in individuals with OCD, revealing increased competitive visuocortical interactions between task cues and task-irrelevant pictures as a function of picture content. However, the magnitude of the visuocortical distraction in the OCD-evoking picture condition was attenuated in comparison to the distraction evoked in the unpleasant picture condition. Subjective valence and arousal ratings also showed considerable differences between the unpleasant and OCD-evoking picture conditions. Previous research examining specific picture categories within the unpleasant IAPS pictures may offer insight into the substantial competition observed in the unpleasant picture condition for the OCD group. Several fMRI studies examining BOLD activation in response to subsets of the unpleasant IAPS pictures in participants with checking, symmetry, harm, and predominantly contamination obsessions revealed unique patterns of activation in response to both disgust-inducing and threat-related pictures (Berlin et al., 2015; Shapira et al., 2003; Stark et al., 2003). In these studies, the disgust-inducing pictures overlapped with the unpleasant pictures used in the present study. Given the considerable impact disgust-inducing pictures have on individuals with OCD, it is possible that the unpleasant picture condition served as a disorder-relevant picture category. The present study is part of a large-scale research initiative investigating obsessive-compulsive anxiety spectrum disorders, and individual picture-level analyses will be conducted once the final sample is collected to examine individual differences in attention capture across specific picture subtypes.

Interestingly, the control analysis indicated attenuated group differences in the magnitude of the visuocortical distraction in response to the pleasant and unpleasant distractor pictures when the OCD-evoking picture condition was excluded from the planned contrast. The results of the control analysis and behavioral analysis reflect the tendency for paradigms presenting pleasant, neutral, and unpleasant IAPS pictures to reliably produce robust visuocortical and behavioral indices of emotional processing (Cuthbert et al., 2000; Keil et al., 2002). However, measures of individual differences obtained from robust, reliable indices of cognition are subject to the “reliability paradox” proposed by Hedge and colleagues (2018). This notion addresses how established cognitive and behavioral tasks may not be suitable for the examination of individual differences due to high reliability of responses elicited in these tasks. To combat the reliability paradox, Haines and colleagues (2025) advocate for use of computational models of cognitive constructs to extract meaningful parameter estimates of neural processes, improving the robustness and reliability of measures obtained from established experimental tasks. Therefore, the present investigation applied the DUC model to further examine the visuocortical trade-off between the task-evoked and picture-evoked responses for each picture condition, and across groups (De Echegaray et al., 2024; Müller et al., 2026). Modeling competitive interactions in visual cortex confirmed the strength of the attentional capture in response to the unpleasant pictures in the OCD group, corroborating the results of the univariate analysis. However, the DUC model also revealed substantial evidence of heightened distraction in the OCD group driven by the OCD-evoking picture condition, which was not readily observed in the univariate analysis.

The present study provides robust evidence of heightened distraction under competition in OCD, largely driven by attentional biases in response to unpleasant and OCD-evoking task-irrelevant distractor pictures. These findings align with prior electrophysiological evidence of enhanced visual processing and attentional selection, particularly in response to aversive and disorder-relevant contexts, which is indicative of hypervigilance associated with obsessive-compulsive psychopathology (Ayomen et al., 2025; Dieterich et al., 2017; Paul et al., 2016).

## Acknowledgements

This research was supported by grant R01MH135426 from the National Institutes of Mental Health awarded to Dr. Carol A. Mathews and Dr. Andreas Keil.

## Declarations

### Ethics approval

All activities related to this research project were completed as a part of a large-scale research initiative investigating obsessive-compulsive anxiety spectrum disorders and approved by the University of Florida International Review Board in accordance with the Declaration of Helsinki.

### Consent to participate

Informed consent was obtained from all individual participants included in the study.

### Consent for publication

Consent for publication was obtained from all individual participants included in the study.

### Availability of data and materials

Behavioral data, electrophysiological data, and limited demographic data are openly available in the project Open Science Framework (OSF) repository [https://osf.io/s73d5/overview]. The supplemental demographic data table (Supplemental Table 1) can also be found in the OSF project repository.

### Code availability

All code and scripts related to data processing and analysis are openly available in the project Open Science Framework (OSF) repository [https://osf.io/s73d5/overview].

### Authors’ contributions

Conceptualization: Katherine J. McCain, Estelle Ayomen, Arash Mirifar, Carol A. Mathews, Andreas Keil. Methodology: Katherine J. McCain, Estelle Ayomen, Arash Mirifar, Carol A. Mathews, Andreas Keil. Software: Katherine J. McCain, Estelle Ayomen, Arash Mirifar, Carol A. Mathews, Andreas Keil. Formal Analysis: Katherine J. McCain, Estelle Ayomen, Arash Mirifar, Carol A. Mathews, Andreas Keil. Investigation: Katherine J. McCain, Estelle Ayomen, Arash Mirifar, Heather Simpson Martin, Dominic Demeterfi, Daniel J. McNeil, Gian DePamphilis, Rami Hatem, Robyn Nelson, Grace Melville, Emma Hammes, Alexandra Lee, Ryan McCarty, Mindy Le, Catherine Paciotti, Patricia Coutinho, Carol A. Mathews, Andreas Keil. Resources: Estelle Ayomen, Arash Mirifar, Heather Simpson Martin, Robyn Nelson, Grace Melville, Carol A. Mathews, Andreas Keil. Data Curation: Katherine J. McCain, Estelle Ayomen, Arash Mirifar, Carol A. Mathews, Andreas Keil. Writing – Original Draft: Katherine J. McCain, Estelle Ayomen, Arash Mirifar, Carol A. Mathews, Andreas Keil. Writing – Review & Editing: Katherine J. McCain, Estelle Ayomen, Arash Mirifar, Heather Simpson Martin, Dominic Demeterfi, Daniel J. McNeil, Gian DePamphilis, Rami Hatem, Robyn Nelson, Grace Melville, Emma Hammes, Alexandra Lee, Ryan McCarty, Mindy Le, Catherine Paciotti, Patricia Coutinho, Carol A. Mathews, Andreas Keil. Visualization: Katherine J. McCain, Estelle Ayomen, Arash Mirifar, Carol A. Mathews, Andreas Keil. Supervision: Carol A. Mathews, Andreas Keil. Funding Acquisition: Carol A. Mathews, Andreas Keil.

### Open Practices Statement

The present study was not preregistered. Behavioral data, electrophysiological data, limited demographic data, and code related to data processing and analysis are openly available in the project Open Science Framework (OSF) repository [https://osf.io/s73d5/overview]. The supplemental demographic data table (Supplemental Table 1) can also be found in the OSF project repository.

## Footnotes

1 Given the significant difference in age distribution between the control group and the OCD group, efforts were taken to evaluate the robustness of the main experimental effects when the groups were more homogenous across age. After excluding participants at or over the age of 65, results from the cluster-based permutation analysis remained robust and mirrored those observed in the full sample. There were no significant group differences in age (t(58) = 1.89, p = .064, *d* = 0.49; OCD: 27.42[9.35], M[SD]; Control: 33.63[15.78], M[SD]) or the other demographic variables (race, ethnicity, and gender) after excluding participants at or over the age of 65.

2 An additional control analysis was completed to examine the within-subject and between-subject effects when the planned contrast weights were identical for each group: -1 (Pleasant), 1 (Neutral), -1 (Unpleasant), 1 (OCD-evoking). Results from the cluster-based permutation analysis remained robust and indicated the presence of the same within-subject and between-subject effects that are reported above.

