## Supplemental Table 1 for "Heightened Distraction under Competition in Obsessive-Compulsive Disorder"

### Supplemental Material

Supplemental Table 1. Demographic data presented as frequency and percentage.

| | | Total (n = 64) | Control Group (n = 31) | | OCD Group (n = 33) | |
|---|---|---|---|---|---|---|
| | | Frequency | Frequency | Percentage | Frequency | Percentage |
| Gender[a] | Female | 49 | 22 | 71.0 | 27 | 81.8 |
| | Male | 14 | 9 | 29.0 | 5 | 15.2 |
| | Prefer not to answer | 1 | 0 | 0.0 | 1 | 3.0 |
| Race[b] | Asian | 3 | 3 | 9.7 | 0 | 0.0 |
| | American Indian/Alaskan Native | 1 | 0 | 0.0 | 1 | 3.0 |
| | Black/African American | 5 | 4 | 12.9 | 1 | 3.0 |
| | Unknown/Prefer not to answer | 1 | 0 | 0.0 | 1 | 3.0 |
| | White/ Caucasian | 54 | 24 | 77.4 | 30 | 91 |
| Ethnicity[c] | Hispanic/Latino | 12 | 5 | 16.1 | 7 | 21.2 |
| | Non-Hispanic/Non-Latino | 52 | 26 | 83.9 | 26 | 78.8 |
| Age[d] | <19 | 5 | 2 | 6.5 | 3 | 9.1 |
| | 20-29 | 34 | 13 | 41.9 | 21 | 63.7 |
| | 30-39 | 9 | 5 | 16.1 | 4 | 12.1 |
| | 40-49 | 4 | 0 | 0.0 | 4 | 12.1 |
| | 50-59 | 6 | 5 | 16.1 | 1 | 3.0 |
| | >60 | 6 | 6 | 19.4 | 0 | 0.0 |
| | Average and standard deviation | | 38.1(19.0) | | 27.4(9.4) | |

[a]Chi-square test showed no significant differences in gender across groups $X^2(2, N = 64) = 2.59$, p = 0.27, Cramer's V = 0.20.

[b]Chi-square test showed no significant differences in race across groups $X^2(4, N = 64) = 7.41$, p = 0.12, Cramer's V = 0.34.

[c]Chi-square test showed no significant differences in ethnicity across groups $X^2(1, N = 64) = 0.27$, p = 0.60, Cramer's V = 0.07.

[d]Independent Welch's t-test showed significant differences in age across groups $t(43.16) = 2.84$, p = .007, $d = 0.72$.
